# Sleep and the gut microbiome: antibiotic-induced depletion of the gut microbiota reduces nocturnal sleep in mice

**DOI:** 10.1101/199075

**Authors:** Jonathan Lendrum, Bradley Seebach, Barrett Klein, Sumei Liu

## Abstract

Several bacterial cell wall components such as peptidoglycan and muramyl peptide are potent inducers of mammalian slow-wave sleep when exogenously administered to freely behaving animals. It has been proposed that the native gut microflora may serve as a quasi-endogenous pool of somnogenic bacterial cell wall products given their quantity and close proximity to the intestinal portal. This proposal suggests that deliberate manipulation of the host's intestinal flora may elicit changes in host sleep behavior. To test this possibility, we evaluated 24 h of sleep-wake behavior after depleting the gut microbiota with a 14 d broad-spectrum antibiotic regimen containing high doses of ampicillin, metronidazole, neomycin, and vancomycin. High-throughput sequencing of the bacterial 16S rDNA gene was used to confirm depletion of fecal bacteria and sleep-wake vigilance states were determined using videosomnography techniques based on previously established behavioral criteria shown to highly correlate with standard polysomnography-based methods. Additionally, considering that germ-free and antibiotic-treated mice have been earlier shown to display increased locomotor activity, and since locomotor activity has been used as a reliable proxy of sleep, we suspected that the elevated locomotor activity previously reported in these animals may reflect an unreported reduction in sleep behavior. To examine this potential relationship, we also quantified locomotor activity on a representative subsample of the same 24 h of video recordings using the automated video-tracking software ANY-maze. We found that antibiotic-induced depletion of the gut microbiota reduced nocturnal sleep, but not diurnal sleep. Likewise, antibiotic-treated mice showed increased nocturnal locomotor activity, but not diurnal locomotor activity. Taken together, these results support a link between the gut microbiome and nocturnal sleep and locomotor physiology in adult mice. Additionally, our findings indicate that antibiotics may be insomnogenic via their ability to diminish gut-derived bacterial somnogens. Given that antibiotics are among the most commonly prescribed drugs in human medicine, these findings have important implications for clinical practice with respect to prolonged antibiotic therapy, insomnia, and other idiopathic sleep-wake and circadian-rhythm disorders affecting an estimated 50-70 million people in the United States alone.

**Highlights:**

- 14 d broad-spectrum antibiotic treatment effectively depletes the gut microbiota.
- Gut microbiota depletion reduces nocturnal sleep, but not diurnal sleep.
- Gut microbiota depletion increases nocturnal locomotion, but not diurnal locomotion.
- Antibiotics may be insomnogenic: implications for idiopathic sleep disorders.

## 1. Introduction

A link between microbes and sleep first became plausible when a potent sleep-inducing substance called “Factor S” was extracted from the brains of sleep-deprived goats (Fencl et al., 1971). Factor S was later identified as muramyl peptide (MP), a unique bacterial cell wall component (BCWC) derived from peptidoglycan (PGN) (Krueger et al., 1982a). In the intervening years, it has become increasingly clear that not only MP, but also PGN and lipopolysaccharide, are potent inducers of mammalian slow-wave sleep (SWS) when exogenously administered to freely-behaving animals (Krueger & Majde, 1994). For instance, just one-hundred picomoles of MP given to rabbits intracerebroventricularly produces a 25% increase in the duration of SWS (individual bouts and total) and a 50% increase in the individual slow-wave voltages over a period of about 6-8 h (Krueger et al., 1982b).

It has been proposed that the commensal gut microflora may serve as a quasi-endogenous pool of these somnogenic BCWCs given their quantity and close proximity to the intestinal portal (Brown, 1995; Krueger et al., 1985). A relationship between coprophagy, bacterial colonization of the neonatal gastrointestinal tract, and the ontogeny of SWS and hematopoiesis has also been proposed (Brown et al., 1988; Korth et al., 1995). If proven correct, similar to essential amino acids and vitamins, BCWCs may be an exogenous requirement for normal physiological processes, in this case for the upkeep of host defense mechanisms and for the generation of SWS (Krueger & Karnovsky, 1995).

Support for the involvement of the gut microbiota (and hence BCWCs) in regulating host sleep behavior comes in part from the findings of Rhee & Kim (1987) that showed a marked decrease in gastrointestinal microflora in psychiatric insomnia patients. Furthermore, several neurological disorders that are associated with sleep disturbances have also been linked to changes in the gut microbiome, including chronic fatigue syndrome (Giloteaux et al., 2016), irritable bowel syndrome (Labus et al., 2017), and obstructive sleep apnea (Durgan, 2017). Notably, Arentsen and colleagues (2017) recently demonstrated that PGN derived from the commensal gut microbiota is translocated from the intestinal mucosa into circulation and across the blood-brain barrier under normal physiological conditions. And perhaps most intriguing, several studies have shown that the gut microbiome rhythmically fluctuates in both community composition and gene expression in a circadian-dependent manner (Leone et al., 2015; Thaiss et al., 2016; Zarrinpar et al., 2014). Taken together, these considerations led to the untested hypothesis that under steady-state conditions, somnogenic BCWCs translocate the gut wall at a basal level contributing to normal sleep regulation, whereas the major enhancements in sleep following exogenous administration of BCWCs are an acute amplification of these physiological processes to compensate for an increased pool of BCWCs over and above the purported endogenous BCWC pool (Krueger & Opp, 2016). However, much work is needed to substantiate and clarify this hypothesis before causality can be established.

The bacterial origin of sleep-inducing substances suggests that deliberate manipulation of the host's intestinal flora may elicit changes in host sleep behavior. Multiple studies have found that germ-free (GF) and antibiotic-induced microbiota depleted (AIMD) mice display increased locomotor activity [reviewed by (Vuong et al., 2017)]. Since locomotor activity (immobility) has been used as a reliable proxy of sleep (Fisher et al., 2012), we hypothesized that the elevated locomotor activity previously reported in these animals may reflect an unreported reduction in sleep behavior. The considerable challenges associated with behavioral testing of GF mice such as cost, labor, and housing restrictions, prompted us to examine whether broad-spectrum antibiotic-induced depletion of the gut microbiota reduces sleep and increases locomotor activity in adult mice. Here, we present an additional line of evidence supporting a role for the gut microbiome in the regulation of nocturnal sleep and locomotor behaviors. We also call attention to the clinically-relevant implications of our finding that antibiotics may be insomnogenic, and briefly speculate as to how this ‘collateral damage’ may be of significance to both practicing physicians and to the millions of people around the world that chronically suffer from insomnia, circadian misalignment, and other idiopathic sleep-wake and circadian-rhythm disorders.

## 2. Methods

### 2.1 Experimental design

Ten genetically inbred C57BL/6 male mice (Harlan Laboratories, Madison, WI) were placed in control (n = 5) or antibiotic-treated (n = 5) groups. Animals were entrained to a standard 12:12 h light:dark cycle in a room isolated from external stimuli with temperatures maintained at 25 ± 1 °C for a minimum of two weeks prior to the experimental protocol. Under these conditions, light onset (0600h) was designated Zeitgeber Time 0 (ZT 0) and dark onset (1800h) as ZT 12. Mice were provided *ad libitum* access to autoclaved water and irradiated food containing the following nutritional formula (in % kcal): 20.5% protein (casein), 69.1% carbohydrates (corn starch, maltodextrin), and 10.4% fat (lard) (Teklad Diet: 08006i, Evigo Bioproducts Inc., Madison, WI). Mice were 14-15 weeks of age at the commencement of experimental treatments. The antibiotic treatment group was orally gavaged once daily for 14 consecutive days with a broad-spectrum antibiotic cocktail consisting of: ampicillin 2.5 mg/ml, metronidazole 2.5 mg/ml, neomycin 2.5 mg/ml, and vancomycin 1.0 mg/ml (Medisca Inc., Plattsburgh, NY). Animals were weighed each morning and a gavage volume of 0.01 ml/g body weight was administered via polypropylene feeding tubes (Instech Laboratories Inc., Plymouth Meeting, PA) without sedation. Fresh antibiotic concoction was prepared daily using autoclaved water as the vehicle. The control group was sham-treated with vehicle by oral gavage at the same volume, frequency, and duration as the antibiotic group. Mice were sacrificed the morning of day 15 via CO_2_ inhalation, followed by exsanguination. All studies were approved by and performed in full accordance with the University of Wisconsin – La Crosse Institutional Animal Care and Use Committee.

### 2.2 Microbiome sequencing

Upon completion of the antibiotic regimen, total DNA was extracted from freshly voided feces using QIAamp DNA Stool Mini Kit (Qiagen Inc., Germantown, MD). The bacterial 16S rDNA gene was amplified with barcoded fusion primers targeting the V3-V4 region and subjected to Illumina (MiSeq) sequencing-by-synthesis as previously described (Klindworth et al., 2013). The QIIME pipeline was used for quality filtering of DNA sequences, demultiplexing, taxonomic assignments, and calculating α and β diversity indices (Caporaso et al., 2012). A dendrogram of bacterial phylogeny was constructed using PhyloT (http://phylot.biobyte.de) and visualized within Interactive Tree of Life (http://itol.embl.de).

### 2.3 Videosomnography

Throughout the duration of the experiment, mice were individually housed in polypropylene cages fitted with video-monitoring equipment capable of recording high-definition video. One miniature, infrared-capable video camera (Sony 138 CMOS with varifocal lens) and one light-emitting-diode (CMVision IR3Wide Angle Array Illuminator) were mounted above each cage to permit continuous recording of animals under light or dark conditions. Cameras were connected to a 16-channel digital video recorder (Dripstone LLC, Los Angeles, California) and set to record at a resolution of 720 x 480 pixels at 30 frames per second. Immediately following completion of the antibiotic regimen, animals were continuously video-recorded for 24 h beginning at 0600 h (ZT 0) on day 14. Cumulative sleep-wake time was visually determined (no attempt was made to discriminate between individual sleep stages) from video recordings using established behavioral criteria previously shown to highly correspond to standard polysomnography-based techniques (Balbir et al., 2008; Storch et al., 2004; Van Twyver et al., 1973). Briefly, mice were considered to be sleeping when there were no gross body movements, except for slight respiratory movements, while displaying typical quiescent sleeping postures (lateral fetal position or prone with head resting on body). Wakefulness was characterized by elevated respiration and the presence of prolonged movement of multiple limbs and/or head (e.g., kicking, stretching, grooming). Sleep fragmentation was scored when the ongoing active or resting phase was interrupted for at least 5 s, or if there were clear and distinct signs of transient awakening. Sleep onset latency was determined for each circadian phase by measuring the time relative to lights – on/lights-off at which each mouse began to sleep (an indicator of circadian regulation of sleep timing), whereas sleep inertia was determined by measuring the time relative to lights – on/lights-off at which each mouse awoke from the principal bout of sleep (an indicator of homeostatic sleep pressure). The diurnal variation of sleep across the entire 24 h was assessed by calculating the total sleep ratio between circadian light and dark phases. Evaluators used Interact (v16) video-annotation software (Mangold International, Arnstorf, Germany) to timestamp sleep-wake behaviors and to compile data.

### 2.4 Locomotor activity

Analysis of locomotor activity was performed on a subsample of video recordings that were representative of each circadian phase using automated video-tracking software as described by Fisher et al. (2012). Here, the program ANY-maze (v5; Stoelting Co., Wood Dale, IL) was used to track and quantify several locomotor activity parameters throughout two 60-min circadian light phase periods (ZT 2-3, ZT 8-9) and two parallel 60-min circadian dark phase periods (ZT 16-17, ZT 22-23). Center (body) position track plots and mean center position occupancy heat maps were generated by the ANY-maze program.

### 2.5 Data analysis

Results were expressed as the mean ± standard error of the mean (SEM), with *n* denoting the number of animals used. Statistical differences between α-diversity indicies were determined by the Mann-Whitney *U* test. Statistical differences between sleep results were determined by the unpaired Student's t test with two-tailed p values less than 0.05 considered significant and reported as *p < 0.05, **p < 0.01. Experimental assignments were randomized, data processing was blind, and statistical analysis was performed using GraphPad Prism version 7.0 (GraphPad Software, La Jolla, CA).

## 3. Results

### 3.1 Broad-spectrum antibiotic treatment effectively depleted the gut microbiota

To verify the antibiotic regimen effectively depleted the gut microbiota, we first examined the structure and diversity of bacteria at a local scale, known as α-diversity. Regardless of the metric employed, α-diversity measures were dramatically decreased by antibiotic treatment. Here, antibiotics reduced the total number of observed species (p < 0.01, Fig. 1A), the rarified species richness (Chao1 index) (p < 0.01, Fig. 1B), and the phylogenetic diversity (PD whole tree index) (p < 0.01, Fig. 1D). The Shannon H index, which takes into account the number of species detected as well as the relative abundance of each bacterial lineage present in the samples, exhibited a similar, but even more pronounced pattern of decreased ecological complexity as a result of prolonged antibiotic treatment (p < 0.01, Fig. 1C). Evident by the leveling (asymptote) of rarefaction curves, the majority of operational taxonomic units (OTUs, i.e. bacterial species) were captured in the α-diversity analysis, which indicates that the small number of shared OTUs was indeed a result of depleting the gut microbiota, and not due to a lack of sequencing depth.

**Figure 1.**
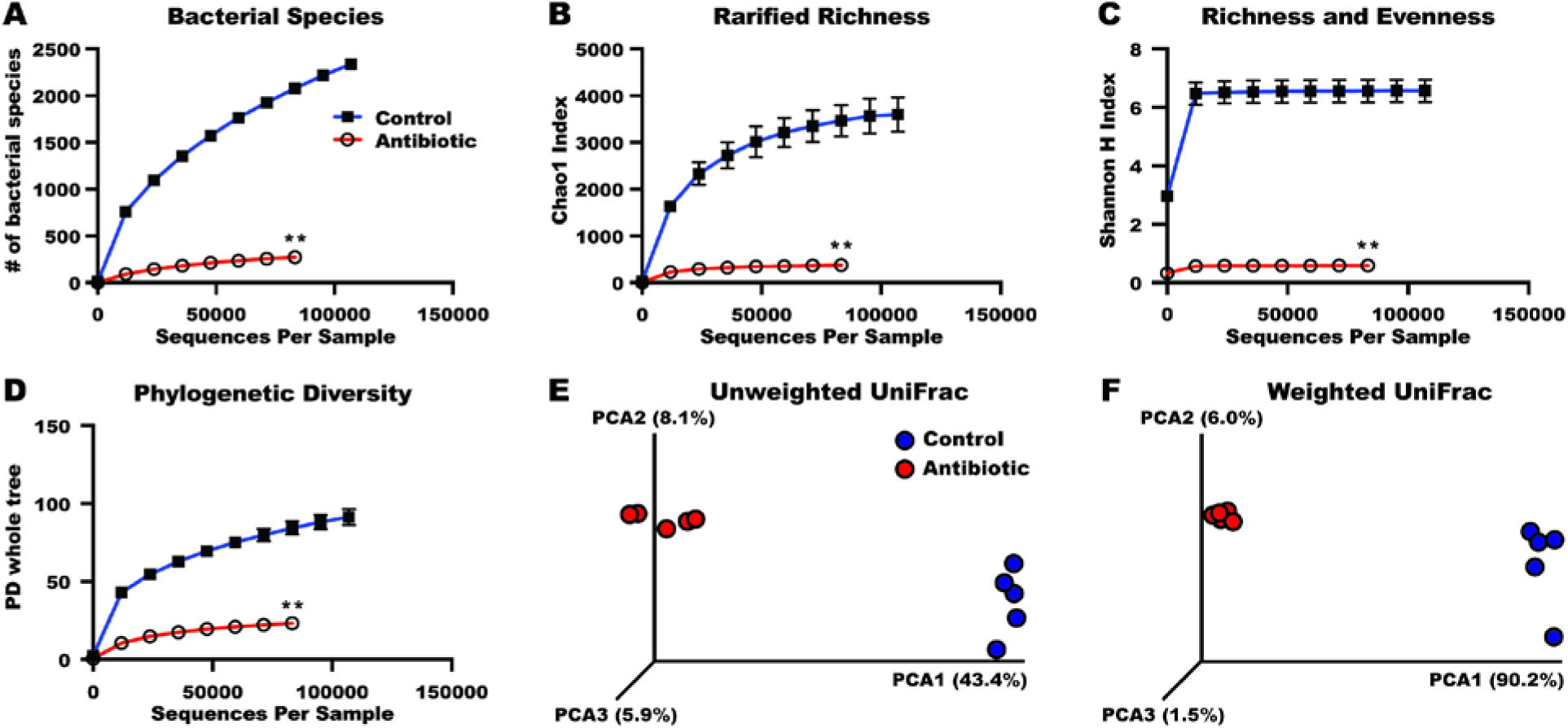
High-throughput sequencing of the bacterial 16S rDNA gene revealed 14 days of broad-spectrum antibiotic treatment via oral gavage effectively depleted fecal bacteria. Upon completion of the antibiotic regimen, total DNA was extracted from freshly voided feces (n = 5 samples per group) and the bacterial 16S rDNA gene was amplified with primers targeting the V3-V4 region and subjected to Illumina (MiSeq) sequencing-by-synthesis. The QIIME pipeline was subsequently used for quality filtering of DNA sequences, assigning taxonomy, and for calculating α **(A-D)** and β-diversity **(EF)** indicies. Antibiotic treatment markedly reduced bacterial α-diversity (within-sample) metrics, including **(A)** the number of bacterial species (measures richness), **(B)** the Chao1 index (measures rarified richness), **(C)** the Shannon H index (measures richness and evenness), and **(D)** the phylogenetic diversity (PD) whole tree index (measures phylogenetic richness). Rarefaction plot error bars represent the 95% confidence interval based on OTUs detected using a sequence similarity threshold of 97%. Statistical significance between groups was determined by the Wilcoxon rank-sum test and reported as **p < 0.01. To compare the microbial community composition among samples, global β-diversity was determined by principal coordinate analysis (PCoA) based on **(E)** unweighted UniFrac distance (uses the presence and absence of OTUs and phylogeny) and **(F)** weighted UniFrac distance (uses the abundance information of OTUs and phylogeny). Each axis represents the percent of total microbial variation between samples with the x-axis representing the greatest dimension of variation and the y and z-axes representing the second and third greatest dimensions of variation, respectively. Samples clustered with respect to antibiotic treatment along all three principal coordinate axes in both the unweighted (emphasizes rare taxa) and weighted (emphasizes abundant taxa) analyses.

We then calculated β-diversity to reveal global scale effects of antibiotics on the full composition of the fecal microbiome. In the unweighted analysis (Fig. 1E), where UniFrac distances were calculated based on the presence/absence of OTUs and phylogeny, the first principal coordinate axis separated samples on the basis of antibiotic treatment and explained 43.4% of the microbial variance. In the weighted analysis, where UniFrac distances were calculated based on abundance information of OTUs and phylogeny, antibiotic-treated samples also clustered along the first coordinate axis (Fig. 1F) which represented 90.2% of the total microbial variance, indicating that most of the predominant commensal bacteria were eradicated by antibiotic treatment. Furthermore, antibiotic-treated samples were tightly grouped in the weighted analysis with respect to the second (6.0%) and third (1.5%) coordinate axes, suggesting that the residual bacteria were more homogeneous than those of control animals. Moreover, the greater separation of sample clusters in the weighted analysis, which emphasizes abundant taxa, compared to the unweighted analysis, which emphasizes rare taxa, indicates that antibiotics diminished the quantity of bacteria more than the diversity.

### 3.2 Depletion of the gut microbiota reduced dark phase sleep, but not light phase sleep

Immediately following completion of the antibiotic regimen, 24 h of sleep-wake behavior was assessed via videosomnography and depicted as a standard bihourly mean percent sleep plot (Fig. 2A). With respect to the circadian light phase, when nocturnal animals are typically inactive, antibiotic-treated mice slept approximately the same as their control counterparts (Fig. 2B). No statistical difference in any sleep parameter was detected during the circadian light phase. However, we found several notable differences between antibiotic and control groups during the circadian dark phase. Here, our principal finding was that mice depleted of their gut microbiota slept 13 ± 3% less than control mice (p < 0.01, Fig. 2C). Reflecting this reduction in dark phase sleep, the average duration of a bout of wakefulness was markedly greater in mice treated with antibiotics (p < 0.05, Supplementary Table 2); yet no statistically significant differences in sleep bout duration or sleep fragmentation were identified. Lastly, antibiotic-treated mice delayed dark-phase sleep onset by more than 20 min compared with control mice (p < 0.05, Supplementary Table 2), and woke later from the principal bout of dark phase sleep (p < 0.05, Supplementary Table 2).

**Figure 2.**
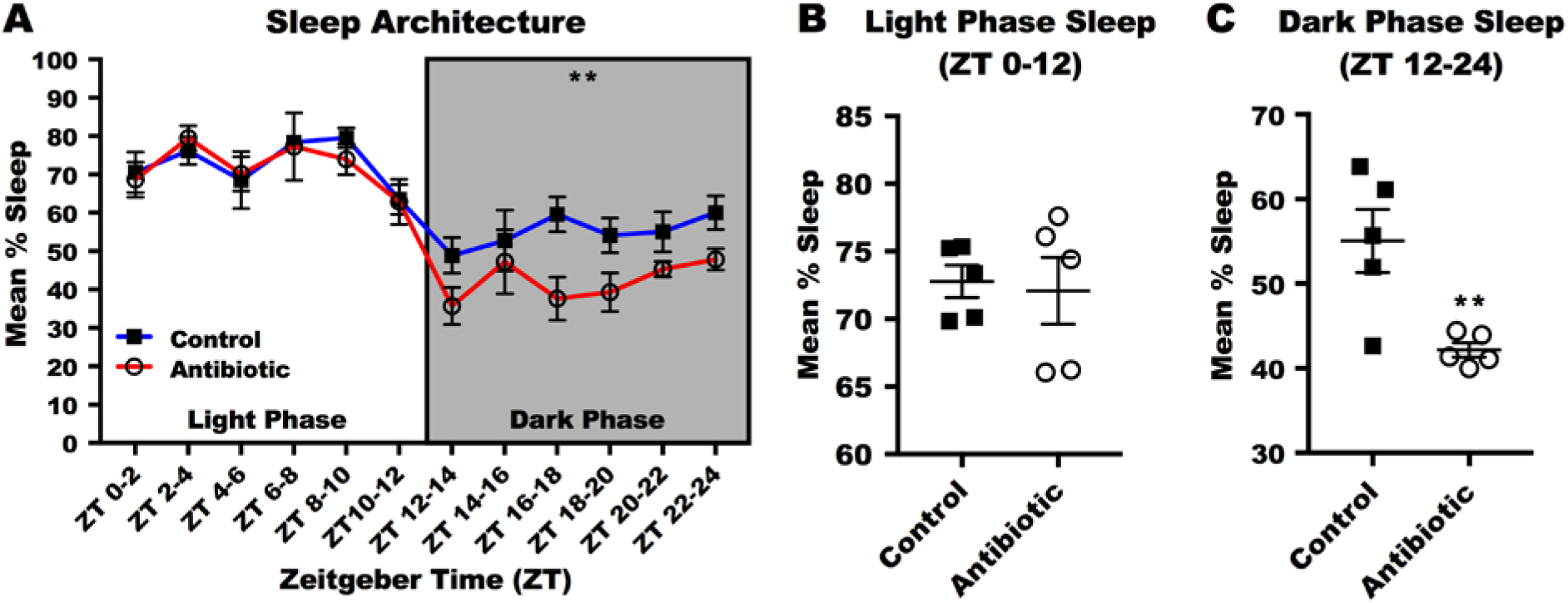
Antibiotic-induced depletion of the gut microbiota reduced dark phase, but not light-phase sleep. Immediately following completion of the antibiotic-depletion regimen animals were video-recorded for 24 h beginning at 0600h (ZT 0). Cumulative sleep-wake time was visually determined from the video recordings using established behavioral criteria previously shown to highly correlate with standard polysomnography-based techniques (no REM vs NREM discrimination was attempted). Sleep scores were compiled into 2 h time bins across the entire 24 h period and depicted as a standard mean percent sleep plot **(A)** with ZT 0-12 (white background) representing the circadian light phase and ZT 12-24 (gray background) representing the circadian dark phase. Mice confirmed to be depleted of their gut microbiota slept for approximately the same duration as controls during the **(B)** circadian light phase (ZT 0-12), but slept significantly less during the **(C)** circadian dark phase (ZT 12-24). Values are group means ± SEM, with n = 5 for each group. Differences between groups were determined for each circadian phase using unpaired Student t test with two-tailed p values less than 0.05 considered significant and expressed as **p < 0.01.

### 3.3 Depletion of the gut microbiota increased dark phase activity, but not light phase activity

Mirroring the effects of antibiotic treatment on sleep described in section 3.2, we observed a similar phase-dependent pattern of locomotor activity. Depicted by the center (body) position track plots in Fig. 3A, antibiotic-treated mice showed greater total distance traveled during dark phase periods (p < 0.05, Fig. 3D), but not light phase periods (Fig. 3C). Likewise, antibiotic-treated mice reached a greater peak velocity during circadian dark phase periods (p < 0.05, Fig. 3E), but not light phase periods (Fig. 3F), and spent more time occupying the center zone of the cage (Fig. 3B).

**Figure 3.**
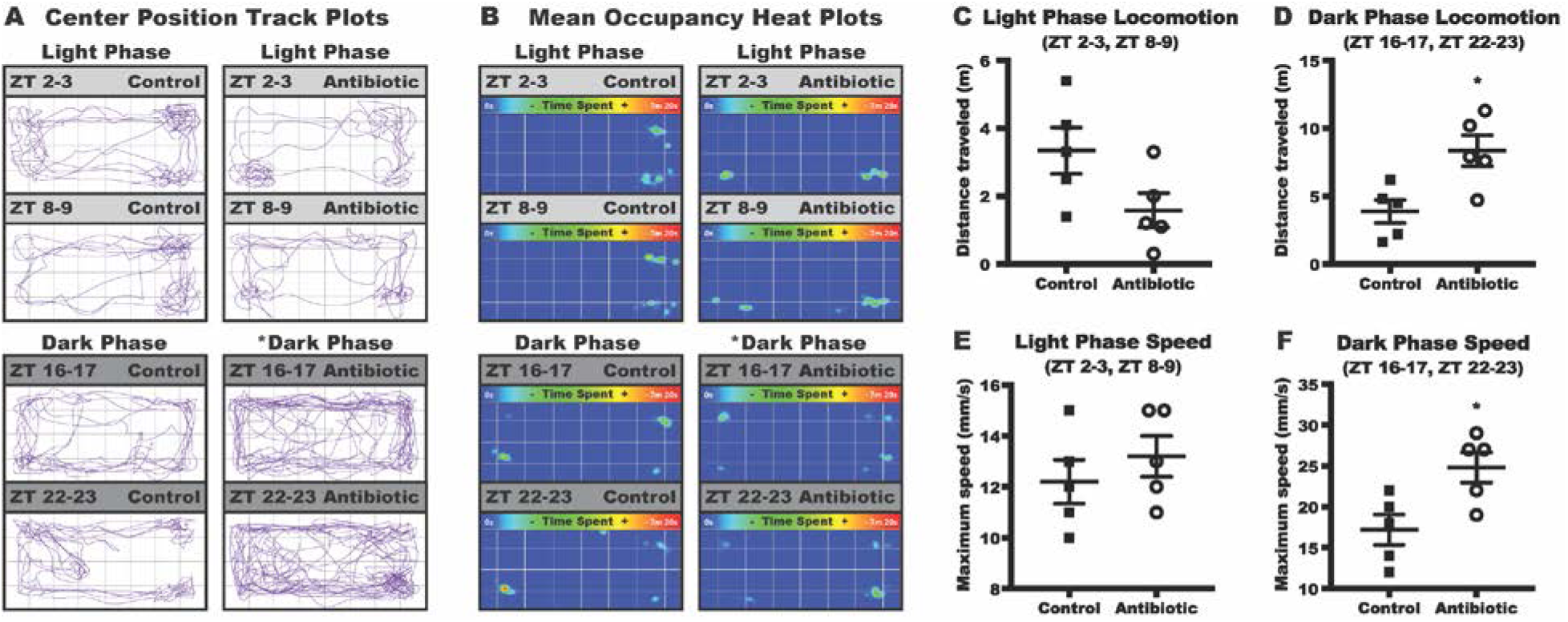
Antibiotic-induced depletion of the gut microbiota increased dark phase, but not light phase locomotor activity. The automated video-tracking software ANY-maze was used to track and quantify several locomotor activity parameters throughout two 60-min circadian light phase periods (ZT 2-3, ZT 8-9) and two parallel 60-min circadian dark phase periods (ZT 16-17, ZT 22-23). **(A)** Combined track plots display the animal's center body position throughout the duration of each 60 min testing periods. **(B)** Mean occupancy heat plots show the animals' tendency to remain primarily in the peripheral cage zone during light and dark-phase testing periods. Antibiotic-treated mice **(C)** traveled approximately the same distance and reached approximately the same **(E)** maximum speed during circadian light phase periods, but traveled a greater distance **(D)** and reached a higher maximum speed **(F)** during circadian dark phase periods. Values are group means ± SEM, with n = 5 for each group. Differences between groups were determined for each circadian phase using unpaired Student t test with two-tailed p values less than 0.05 considered significant and expressed as *p < 0.05, **p < 0.01.

## 4. Discussion

### 4.1 Main findings

In the present study, we show that depleting the gut microbiota with antibiotics reduces circadian dark phase sleep in adult mice. Our data suggest that a reduction of intestinal bacteria decreases dark phase sleep in part by diminishing a potential source of sleep-promoting substances, namely the somnogenic BCWCs. Since diminished sleep correlated with hyperlocomotion, and these effects were confined to the circadian dark phase, it appears that this manipulation specifically interferes with the nocturnal sleep induction process rather than the whole sleep topography.

### 4.2 Antibiotics and behavior

The possibility that these behavioral changes are a consequence of the direct actions of antibiotic treatment *per se* cannot be entirely ruled out because antibiotics are known to have toxic effects on host mitochondrial and ribosomal function (Morgun et al., 2015). However, several observations and inferences strongly suggest that our data are not due to some nonspecific actions of antibiotics. First, if there were any toxic effects due to the treatment we would have expected global changes in locomotor and sleep topography to have occurred, rather than the specific phase-dependent changes seen here. Second, the effects of antibiotic treatment on motor activity and anxiolytic behavior are not likely to be caused by host toxicity since identical behavioral traits are reported in GF mice (Arentsen et al., 2015; Diaz Heijtz et al., 2011; Nishino et al., 2013). Third, GF animals treated with antibiotics show no additional changes in behavior (Bercik et al., 2011; Cox et al., 2014). And perhaps most compelling, Frohlich and colleagues (2016) demonstrated that the same antibiotics used in this study had little to no oral bioavailability and did not enter the brains of mice, thereby providing strong evidence that the AIMD mouse model is a valid and reliable approach to establish causality in gut microbiome-brain/behavior interactions, relative to GF mice.

### 4.3 Potential mechanisms underlying gut microbiome regulation of sleep

The somnogenic properties of BCWCs are thought to be mediated in part by their ability to stimulate Type 1 helper T cells (T_H_1) cells to produce the pro-inflammatory cytokine interleukin (IL)-1β (Dunn, 2006; Opp & Imeri, 1999; Opp & Krueger, 2017). When administered alone, IL-1 also produces large amounts of SWS (Krueger et al., 1984), and pre-treatment with inhibitors of IL-1 attenuates the enhancement of SWS following administration of BCWCs (Imeri et al., 1993), as would be expected if the effector molecule of the immunogenic and somnogenic actions of BCWCs is the cytokine IL-1.

Consequently, given that high exposure to BCWCs stimulates production of pro-inflammatory T_H_1-mediated cytokines that increase sleep, it stands to reason that low exposure to BCWCs (i.e. GF/AIMD mice) may instead encourage the default production of anti-inflammatory T_H_2-mediated cytokines that inhibit sleep (Rogers & Croft, 1999). Indeed, the GF/AIMD immune response is suppressed and T_H_2 biased (Green-Johnson, 2012; Josefsdottir et al., 2017; Mazmanian et al., 2005; Oyama et al., 2001), and anti-inflammatory cytokines have been earlier reported to inhibit spontaneous sleep in freely behaving animals (Krueger et al., 2001; Kushikata et al., 1998, 1999). Taken together, it appears as though one mechanism by which the gut microbiota may regulate normal sleeping patterns is by directing the T_H_1/T_H_2 balance of pro-inflammatory (somnogenic) and anti-inflammatory (insomnogenic) cytokine production (Krueger et al., 1994; Zielinski & Krueger, 2011). As such, we suspect that depleting the antigenic load of BCWCs may have resulted in a diminished basal rate of phagocyte and T_H_1 cell activation, thereby suppressing steady-state production of IL-1, and in turn reducing sleep.

However, cytokines alone do not explain why antibiotics specifically reduced dark phase, but not light phase sleep. We propose that this conspicuous circadian-dependent effect can be explained as a consequence of diminished interactions between BCWCs, IL-1, and the central serotonergic (5-HT) system. To shed light on this feature, we first reflect upon how 5-HT is involved with the regulation of SWS in general, and then we briefly consider how BCWCs and IL-1 interact with 5-HT to induce SWS in an analogous circadian-dependent fashion.

With respect to 5-HT and SWS, data obtained from rats and mice indicates that when 5-HT release is enhanced through the administration of a 5-HT precursor (5-HTP), an increase of SWS is observed specifically during the dark phase of the light-dark cycle, irrespective of whether 5-HTP is given at the beginning of the light or dark phase (Imeri & Opp, 2009; Morrow et al., 2008). With that in mind, now consider the following three assertions regarding how BCWCs and IL-1 interact with 5-HT to induce SWS. First, the SWS induced by BCWCs is accompanied by an increase in 5-HT turnover in several sleep-linked regions of the brain (Masek et al., 1973), and this effect is abolished by electrolytic lesions of raphe serotonergic structures (Masek et al., 1975, 1978). Second, both BCWCs and IL-1 are effective at inducing SWS when administered during the dark phase (Inoue et al., 1984; Meltzer et al., 1989), but are unable to induce SWS when given during the light phase (Fornal et al., 1984; Lancel et al., 1996). Third, 5-HT is essential for BCWCs and IL-1 to exert their effects on sleep because pretreatment with a 5-HT synthesis blocker or antagonism of 5-HT_2_ receptors completely blocks the dark phase SWS induced by BCWCs or IL-1 (Imeri et al., 1997, 1999).

Since the somnogenic effects of 5-HT on SWS are confined to the dark phase, and because the somnogenic effects of BCWCs and IL-1 are also restricted to the dark phase and are accompanied by elevated 5-HT turnover in the brain, combined with the fact that BCWCs and IL-1 are unable to induce SWS when given to animals pre-treated with a 5-HT synthesis blocker, suggests that such dark phase-specific sleep responses reflect a fundamental property of the 5-HT system and thus are likely to be involved with the physiological regulation of sleep by the gut microbiome.

Remarkably, one of the somnogenic BCWCs, namely MP, has been independently shown to be a molecular mimic of 5-HT (Polanski et al., 1992; Ševč ík & Ma šek, 1999). This revelation suggests even a direct involvement of MP in the induction of SWS, possibly by functioning as a 5HT-like neurotransmitter (Root-Bernstein & Westall, 1988). Moreover, since BCWCs such as MP and PGN have been detected in brain tissue (Arentsen et al., 2017; Karnovsky, 1986), and considering that during sleep deprivation MP accumulates in the brain (Pappenheimer et al., 1975), it has to be inferred that there is a physiological low-dose passage of these constituents into brain tissue during normal waking states. It's tempting to speculate that the wake state-associated ingestion of BCWCs (during active feeding) and the subsequent accumulation of BCWCs and IL-1 in the CNS leads beyond a certain threshold to fatigue and sleepiness, finally inducing sleep, during which the clearance of these BCWCs from the CNS is performed (Abstinta et al., 2017; Xie et al., 2013) and the T_H_1/T_H_2 cytokine balance is restituted (Besedovsky et al., 2012). In keeping with this view, the protracted bouts of wakefulness that antibiotic-treated mice displayed here may reflect a diminished rate of BCWC accumulation in the brain over a given waking period, thus lowering the homeostatic sleep pressure threshold and allowing the bout of wakefulness to continue for an extended length of time. In keeping with this speculation, Arentsen and colleagues (2017) found that PGN levels were substantially lower in the brains of GF and AIMD mice compared to mice with an intact gut microbiota. Regardless of whether our sleep results were a reflection of abolishing direct (MP → 5-HT → SWS) or indirect (MP → IL-1 → 5-HT → SWS) interactions, earlier studies using GF and AIMD mice have established that the gut microbiome does in fact regulate host tryptophan and serotonin availability, synthesis, and metabolism both peripherally and centrally under steady-state conditions (Clarke et al., 2013; O'Mahony et al., 2014; Yano et al., 2015), providing further credence to a link between gut microbes, BCWCs, cytokines, 5-HT, and physiological sleep. For these reasons, interactions between the gut microbiome, the immune system, and the 5-HT system appear to be an especially promising target for future investigation.

However, the gut microbiome is a dynamic, living entity capable of producing a vast array of biologically active metabolites that participate in numerous physiological processes. Consequently, there is almost certainly multiple other ways by which the gut microbiome contributes to physiological sleep. For instance, there is a significant body of evidence indicating that the gut microbiota influences the development and later function of the stress system (Foster et al., 2016; Rea et al., 2015). Both GF and AIMD mice have been shown to exhibit HPA axis hyper-responsivity as evident by a nearly 3-fold increase in blood corticosterone levels (Desbonnet et al., 2015; Frohlich et al., 2016; Mukherji et al., 2013; Sudo et al., 2004). Because corticosterone is both a powerful immunosuppressant (Kainuma et al., 2009) as well as a well-known arousal-promoting substance (Vazquez-Palacios et al., 2001), it's conceivable that the reduced sleep and elevated locomotor activity observed in this study, as well as the elevated locomotor activity and immunosuppression previously recognized in GF/AIMD animals, may also reflect a heightened basal state of stress. Clearly much work is needed to establish or debunk these potential relationships and also to disentangling them from one another with respect to their relative contribution to homeostatic sleep regulation.

### 4.4 Study limitations and future research directions

One clear limitation of the present study is the inability of current videosomnography techniques to discriminate between specific sleep stages. It's promising that this limitation may soon be overcome as several groups have begun developing high-throughput behavioral monitoring systems using infrared beams (Pack et al., 2007), piezoelectric sensors (Yaghoby et al., 2016), and sensitive videosomnography algorithms (McShane et al., 2012) that allow for the non-invasive analysis of individual sleep stages. Given that BCWCs are known to specifically increase SWS, we suspect that the reduction in nocturnal sleep observed here may be a reflection of diminished SWS time, but this remains unsubstantiated. Moreover, our finding of a correlation between reduced sleep and hyperactivity appears to support our secondary hypothesis that the elevated locomotor activity previously documented in GF/AIMD mice is a consequence of diminished sleep; however, given our small sample size it will be important to corroborate this by finally characterizing sleep in germ-free rodents. We speculate that the only reason why sleep has not already been characterized in GF rodents is due to the invasive nature of polysomnography (Kis et al., 2014) combined with the fact that behavioral testing of GF animals is restricted due to required housing of the mice in sterile isolator units to maintain microbiologically-free conditions (Al-Asmakh & Zadjali, 2015). Although logistically difficult given that GF animals will be colonized with bacteria shortly after their removal from the GF unit (Hansen et al., 2015), it's conceivable (and highly advisable) that future researchers could overcome this issue by removing the animals and performing the electrode implantation surgery and subsequent polysomnography testing in a sterile room to avoid contamination. Alternatively, this difficulty could be circumvented altogether by utilizing non-invasive telemetry or videosomnography-based techniques like those used in the current study.

Several important questions remain unanswered. Given that the gut microbiome has been compared to an enteric “fingerprint” due to the large degree of specificity to the individual (Consortium, 2013), a logical question arises: to what extent is the inter-individual variation in sleep duration and timing a reflection of inter-individual variation in gut microbiome composition and structure? Considering that BCWCs increase SWS in a dose-dependent manner (Meltzer et al., 1989), taken together with our results that depletion of the gut microbiota (and hence BCWCs) reduces sleep, seems to suggest that the amount of non-specific bacterial exposure may be the primary driving factor for normal sleep and immune regulation, whereas the diversity of bacteria and timing of exposure may be secondary to this effect. Assuming that sleep of GF animals is eventually characterized, this should be testable fairly easily by administering gnotobiotic cultures containing equal (and unequal) quantities of phylogenetically similar (and dissimilar) bacterial communities. Alternatively, this could be tested by treating conventional animals with narrow-spectrum antibiotics that kill specific lineages of bacteria while leaving others intact.

Heeding the vast array of metabolites produced by the gut microbiome, it will be important to distinguish the relative contribution of BCWCs in relation to other biologically active bacterial products such as short-chain fatty acids, and to separate these factors from other environmental factors, including diet and host genetics, in order to accurately characterize the influence of these key microbial products on physiological sleep. To do so, one may consider inoculating GF animals with one of the atypical strains of bacteria that do not have a cell wall (i.e. *Mycoplasma)* and therefore do not yield somnogenic BCWCs, but still allow for a viable bacterial load in the gut. Furthermore, given that some disturbances to the gut microbiome have been found to cause permanent changes to brain function and behavior (Cox et al., 2014; Desbonnet et al., 2015; Diaz Heijtz et al., 2011), future studies should investigate whether reconstituting the gut microbiome via probiotic administration or fecal transplant could return sleep to pre-antibiotic treatment levels. Lastly, due to the recent discovery of the glymphatic system (Jessen et al., 2015) and meningeal lymphatic vasculature (Absinta et al., 2017; Aspelund et al., 2015), it may also be advantageous to characterize the temporal features and dissemination pathways of BCWCs entry into (Louveau et al., 2015) and clearance out of the brain (Raper et al., 2016) with respect to both monophasic and polyphasic as well as to nocturnal and diurnal sleepers.

### 4.5 Clinical implications

The Centers for Disease Control and Prevention has deemed insufficient sleep a major public health problem linked to motor vehicle crashes, military casualties, medical and other costly occupational errors that impose a financial burden in excess of $400 billion a year in the United States alone (Colten et al., 2006; Dement & Gelb, 1993; Reynolds et al., 2017). In the present study, we highlight a previously unrecognized risk factor, namely antibiotic-induced dysbiosis, and call attention to the profound implications for clinical practice given that more than 270 million antibiotic prescriptions are issued in the US each year (Suda et al., 2014). There is now direct experimental evidence in mice (this study), rats (Brown et al., 1990), and humans (Nonaka et al., 1983) indicating that antibiotics may be insomnogenic. Additionally, Perlis et al. (2006) used the Physicians' Desk Reference to document the clinical occurrence of insomnia with respect to seven classes of antibiotics and found that five of the classes were associated with insomnia as a side effect. Furthermore, a retrospective analysis of clinical patients identified antibiotics as one category of medications that may cause leukopenia (Shuman et al., 2012). Hence, there is sufficient evidence to suggest that antibiotics are not benign with respect to sleep and that antibiotic-induced perturbation of the gut microbiota and ensuing immunosuppression may serve as a precipitating factor in the pathogenesis of insomnia.

Since enhanced sleep appears to be an adaptive process that aids in recuperation and elimination of infections (Opp, 2009; Toth et al., 1993), antibiotic therapy, although helpful with respect to its bactericidal actions, may paradoxically potentiate the infectious process by inhibiting and fragmenting sleep, or instead, by way of Jarisch-Hershimer reaction (Kaplanski et al., 1998). It may be possible to attenuate these adverse effects by co-administering antibiotics with pre/probiotics or bacterial ligands as several preliminary studies have demonstrated improved remission in these patients (Boyanova & Mitov, 2012). However, given that the ontogeny of SWS parallels the colonization of the intestinal tract (Davenne & Krueger, 1987; Greenhalgh et al., 2016; Jouvet-Mounier et al., 1969), and old age is associated with reduced and fragmented sleep as well as dysbiosis (Ingiosi et al., 2013; Langille et al., 2014; Wimmer et al., 2013), the use of antibiotics in these particularly vulnerable populations, especially those with underlying sleep pathologies, should be applied with great caution. Alternatively, whether antibiotic treatment will translate into a useful therapy for idiopathic hypersomnolence and chronic fatigue syndrome, or conversely, whether strategically timed consumption of probiotics will translate into an effective and safe class of sleep sedatives, remains to be seen. Promisingly, Kumar and colleagues (2012) showed that antibiotic pretreatment attenuated chronic fatigue-induced oxidative stress in mice, and Thompson and colleagues (2017) recently demonstrated that dietary prebiotics attenuated stress-induced alterations of sleep in rats, thus giving hope for future studies in this direction.

## 5. Conclusion

In conclusion, it seems quite likely that the gradual breakdown of intestinal flora increases circulating BCWCs which serve as immune adjuvants, serotonin mimics, and physiological regulators of mammalian sleep. Our findings highlight a unique microbiome feature, and given its targetable nature, have important implications for preventative care as well as early diagnosis and therapeutic intervention in the management of sleep-wake and circadian rhythm disorders, the occurrence of which will inevitably rise as technological advances continue to pressure the limits of human adaptability.

## 6. Data Access Statement

Supplementary data associated with this article can be found in the online version, at https://www.biorxiv.org/.

## 7. Authors' Contributions

Designed experiments: JL, BS, BK, SL. Performed experiments: JL, BS, SL. Analyzed data: JL, SL. Edited manuscript: JL, BS, BK, SL.

## 9. Acknowledgements

This work was supported by NIH grant R15 DK097460-01A1 (SL) and University of Wisconsin-La Crosse Undergraduate Research and Creativity grants 4-15 and 11-15 (JL). The authors thank the University of Wisconsin-Biotechnology Center DNA Sequencing Facility for providing sequencing and bioinformatics services, Amy Cooper for assisting with animal care duties, and Ronald Zaleski, Dotty Storch, and Christopher Forehand for their time scoring video recordings. JL is indebted to Julian Strucksheats as well as Georgia Bailey and Kip Sullivan for their continued support and advice.

## References

Absinta, M., Ha, S., Nair, G., Sati, P., Luciano, N., Palisoc, M., Louveau, A., Zaghloul, K., Pittaluga, S., Kipnis, J., & Reich, D. (2017). Human and nonhuman primate meninges harbor lymphatic vessels that can be visualized noninvasively by MRI. eLife. 6, e29738. https://doi.org/10.7554/eLife.29738

Al-Asmakh, M., & Zadjali, F. (2015). Use of germ-free animal models in microbiota-related research. J. Microbiol. Biotechnol. 25, 1583–1588. http://dx.doi.org/10.4014/jmb.1501.01039

Arentsen, T., Qian, Y., Gkotzis, S., Femenia, T., Wang, T., Udekwu, K., Forssberg, H., & Diaz Heijtz, R. (2017). The bacterial peptidoglycan-sensing molecule Pglyrp2 modulates brain development and behavior. Mol. Psychiatry. 22, 257–266. https://doi.org10.1038/mp.2016.182

Arentsen, T., Raith, H., Qian, Y., Forssberg, H., & Diaz Heijtz, R. (2015). Host microbiota modulates development of social preference in mice. Microb. Ecol. Health. Dis. 26, 29719. http://dx.doi.orgHYPERLINK “http://dx.doi.org/10.3402/mehd.v26.29719” /10.3402/mehd.v26.29719

Aspelund, A., Antila, S., Proulx, S., Karlsen, T., Karaman, S., Detmar, M., Wiig, H., & Alitalo, K. (2015). A dural lymphatic vascular system that drains brain interstitial fluid and macromolecules. J. Exp. Med. 212, 991–999. https://doi.org/10.1084/ jem.20142290

Balbir, A., Lande, B., Fitzgerald, R., Polotsky, V., Mitzner, W., & Shirahata, M. (2008). Behavioral and respiratory characteristics during sleep in neonatal DBA/2J and A/J mice. Brain. Res. 1241, 84–91. https://doi.org/10.1016/j.brainres.2008.09.008

Bercik, P., Denou, E., Collins, J., Jackson, W., Lu, J., Jury, J., Deng, Y., Blennerhassett, P., Macri, J., McCoy, K., Verdu, E., & Collins, S. (2011). The intestinal microbiota affect central levels of brain-derived neurotropic factor and behavior in mice. Gastroenterol. 141, 599–609. https://doi.org/10.1053/j.gastro.2011.04.052

Besedovsky, L., Lange, T., & Born, J. (2012). Sleep and immune function. Pflugers. Arch. 463, 121–137. https://doi.org/10.1007/s00424-011-1044-0

Boyanova, L., & Mitov, I. (2012). Coadministration of probiotics with antibiotics: why, when and for how long? Expert. Rev. Anti. Infect. Ther. 10. 407–409. http://dx.doi.org/10.1586/eri.12.24

Brown, R. (1995). Muramyl peptides and the functions of sleep. Behav. Brain. Res. 69, 85–90. https://doi.org/10.1016/0166-4328(95)00004-D

Brown, R., Price, R., King, M., & Husband, A. (1988). Autochthonous intestinal bacteria and coprophagy: a possible contribution to the ontogeny and rhythmicity of slow wave sleep in mammals. Med. Hypotheses. 26, 171–175. https://doi.org/10.1016 /0306-9877(88)90096-5

Brown, R., Price, R., King, M., & Husband, A. (1990). Are Antibiotic Effects on Sleep Behavior in the Rat Due to Modulation of Gut Bacteria? Physiol. Behav. 48, 561–565. https://doi.org/10.1016/0031-9384(90)90300-S

Caporaso, J., Lauber, C., Walters, W., Berg-Lyons, D., Huntley, J., Fierer, N., Owens, S., Betley, J., Fraser, L., Bauer, M., Gormley, N., Gilbert, J., Smith, G., & Knight, R. (2012). Ultra-high-throughput microbial community analysis on the Illumina HiSeq and MiSeq platforms. ISME J. 6, 1621–1624. https://doi.org/10.1038/HYPERLINK “https://doi.org/10.1038/ismej.2012.8” ismej.2012.8

Clarke, G., Grenham, S., Scully, P., Fitzgerald, P., Moloney, R., Shanahan, F., Dinan, T., & Cryan, J. (2013). The microbiome-gut-brain axis during early life regulates the hippocampal serotonergic system in a sex-dependent manner. Mol. Psychiatry. 18, 666–673. https://doi.orgorg/10.1038/mp.2012.77” /10.1038/mp.2012.77

Colten, H., & Altevogt, B. (2006). Sleep disorders and sleep deprivation: an unmet public health problem. Washington, DC: The National Academies Press. https://doi.org/10.17226/11617

Consortium, T. H. M. P. (2013). Structure, Function and Diversity of the Healthy Human Microbiome. Nature. 486, 207–214. https://doi.org/10.038/nature1234

Cox, L., Yamanishi, S., Sohn, J., Alekseyenko, A., Leung, J., Cho, I., Kim, S., Li, H., Gao, Z., Mahana, D., Zarate Rodriguez, J., Rogers, A., Robine, N., Loke, P., & Blaser, M. (2014). Altering the Intestinal Microbiota during a Critical Developmental Window Has Lasting Metabolic Consequences. Cell. 158, 705–721. https://doi.org/10.1016/j.cell.2014.05.052

Davenne, D., & Krueger, J. (1987). Enhancement of quiet sleep in rabbit neonates by muramyl dipeptide. Am. J. Physiol. Regul. Integr. Comp. Physiol. 253, R646–654.

Dement, W., & Gelb, M. (1993). Somnolence: its importance in society. Clin. Neurophysiol. 23, 5–14. https://doi.org/10.1016/S0987-7053(05)80278-3

Desbonnet, L., Clarke, G., Traplin, A., O’Sullivan, O., Crispie, F., Moloney, R., Cotter, P., Dinan, T., & Cryan, J. (2015). Gut microbiota depletion from early adolescence in mice: Implications for brain and behaviour. Brain. Behav. Immun. 48, 165–173. https://doi.org/10.1016/j.bbi.2015.04.004

Diaz Heijtz, R., Wang, S., Anuar, F., Qian, Y., Björkholm, B., Samuelsson, A., Hibberd, M., Forssberg, H., & Pettersson, S. (2011). Normal gut microbiota modulates brain development and behavior. Proc. Natl. Acad. Sci. U.S.A. 108, 3047–52. https://doi.org/10.1073/pnas.1010529108

Dunn, A. (2006). Effects of cytokines and infections on brain neurochemistry. Clin. Neurosci. Res. 6, 52–68. https://doi.org/10.1016/j.cnr.2006.04.002

Durgan, D., (2017). Obstructive sleep apnea-induced hypertension: role of the gut microbiota. Curr. Hypertens. Rep. 19, 35. https://doi.org/10.1007/s11906-017-0732-3

Fencl, V., Koski, G., & Pappenheimer, J. (1971). Factors in cerebrospinal fluid from goats that affect sleep and activity in rats. J. Physiol. 216, 565–89. https://doi.org/10.1113/jphysiol.1971.sp00954

Fisher, S., Godinho, S., Pothecary, C., Hankins, M., Foster, R., & Peirson, S. (2012). Rapid assessment of sleep-wake behavior in mice. J. Biol. Rhythms., 27, 48–58. https://doi.org/10.1177/0748730411431550

Fornal, C., Markus, R., & Radulovacki, M. (1984). Muramyl dipeptide does not induce slow-wave sleep or fever in rats. Peptides. 5, 91–95. https://doi.org/10.1016/0196-9781(84)90057-3-9781(84)90057-3

Foster, J., Rinaman, L., & Cryan, J. (2017). Stress & the gut-brain axis: regulation by the microbiome. Neurobiol. Stress. 7, 1–13. https://doi.org/10.1016/j.ynstr.2017.03.001

Frohlich, E., Farzi, A., Mayerhofer, R., Reichmann, F., Jacan, A., Wagner, B., Zinser, E., Bordag, N., Magnes, C., Frohlich, E., Kashofer, K., Gorkiewicz, G., & Holzer, P. (2016). Cognitive impairment by antibiotic-induced gut dysbiosis: analysis of gut microbiota-brain communication. Brain. Behav. Immun. 56, 140–155. https://doi.org/10.1016/j.bbi.2016.02.020

Giloteaux, L., Goodrich, J., Walters, W., Levine, S., Ley, R., & Hanson, M. (2016). Reduced diversity and altered composition of the gut microbiome in individuals with myalgic encephalomyelitis/chronic fatigue syndrome. Microbiome. 4, 30. https://doi.org/10.1186/s40168-0171-4

Greenhalgh, K., Meyer, K., Aagaard, K., & Wilmes, P. (2016). The human gut microbiome in health: establishment and resilience of microbiota over a lifetime. Environ. Microbiol. 18, 2103–2116. https://doi.org/10.1111/1462-2920.13318

Green-Johnson, J. (2012). Immunological Responses to Gut Bacteria. J. AOAC Int. 95, 35–49. https://doi.org/10.5740/jaoacint.SGE_Green-Johnson

Hansen, A., Krych, Ł., Nielsen, D., & Hansen, C. (2015). A review of applied aspects of dealing with gut microbiota impact on rodent models. ILAR. J. 56, 250–264. https://doi.org/10.1093/ilar/ilv010

Imeri, L., & Opp, M. (2009). How (and why) the immune system makes us sleep. Nature. Rev. Neurosci. 10, 199–210. https://dx.doi.org/10.1038/nrn2576

Imeri, L., Bianchi, S., & Mancia, M. (1997). Muramyl dipeptide and IL-1 effects on sleep and brain temperature after inhibition of serotonin synthesis. Am. J. Physiol. Regul. Integr. Comp. Physiol. 273, R1663–R1668.

Imeri, L., Mancia, M., & Opp, M. (1999). Blockade of 5-hydroxytryptamine (serotonin)-2 receptors alters interleukin-1-induced changes in rat sleep. Neuroscience. 92, 745–749. https://doi.org/10.1016/S0306-4522(99)00006-8

Imeri, L., Opp, M., & Krueger, J. (1993). An IL-1 receptor and an IL-1 receptor antagonist attenuate muramyl dipeptide-and IL-1-induced sleep and fever. Am. J. Physiol. 265, 907–913.

Ingiosi, A., Opp, M., & Krueger, J. (2013). Sleep and immune function: glial contributions and consequences of aging. Curr. Opin. Neurobiol. 23, 806–811. https://doi.org/10.1016/j.conb.2013.02.003

Inoue, S., Honda, K., Komoda, Y., Uchizono, K., Ueno, R., & Hayaishi, O. (1984). Differential sleep-promoting effects of five sleep substances nocturnally infused in unrestrained rats. Proc. Natl. Acad. Sci. U.S.A. 81, 6240–6244. https://doi.org/10.1073/pnas.81.19.6240

Jessen, N., Munk, A., Lundgaard, I., & Nedergaard, M. (2015). The Glymphatic System: A Beginner’s Guide. Neurochem. Res. 40, 2583–2599. https://doi.org/10.1007 /s11064-015-1581-6

Josefsdottir, K., Baldridge, M., Kadmon, C., & King, K. (2017). Antibiotics impair murine hematopoiesis by depleting the intestinal microbiota. Blood. 129, 729–739. https://doi.org/10.1182/blood-2016-03-708594

Jouvet-Mounier, D., Astic, L., & Lacote, D. (1969). Ontogenesis of the states of sleep in rat, cat, and guinea pig during the first postnatal month. Dev. Psychobiol. 2, 216–239. https://doi.org/10.1002/dev.420020407

Kainuma, E., Watanabe, M., Tomiyama-Miyaji, C., Inoue, M., Kuwano, Y., Ren, H., & Abo, T. (2009). Association of glucocorticoid with stress-induced modulation of body temperature, blood glucose and innate immunity. Psychoneuroendocrinology. 34, 1459–1468. https://doi.org/10.1016/j.psyneuen.2009.04.021/j.psyneuen.2009.04.021

Kaplanski, G., Granel, B., Vaz, T., & Durand, J. (1998). Jarisch-Herxheimer reaction complicating the treatment of chronic Q fever endocarditis: elevated TNFα and IL-6 serum levels. J. Infect. 37, 83–84. https://doi.org/10.1016/S01634453(98)91120-3

Karnovsky, M. (1986). Muramyl peptides in mammalian tissues and their effects at the cellular level. Fed. Proc. 45, 2556–2560.

Kis, A., Szakadot, S., Kovacs, E., Gocsi, M., Simor, P., Gombos, F., Topal, J., Miklosi, A., & Bodizs, R. (2014). Development of a non-invasive polysomnography technique for dogs (Canis familiaris). Physiol. Behav. 130, 149–156. https://doi.org/10.1016/j.physbeh.2014.04.004

Klindworth, A., Pruesse, E., Schweer, T., Peplies, J., Quast, C., Horn, M., & Glöckner, F. (2013). Evaluation of general 16S ribosomal RNA gene PCR primers for classical and next-generation sequencing-based diversity studies. Nucleic. Acids. Res. 41, 1–11. https://doi.org/10.1093/nar/gks808/nar/gks808

Korth, C., Wehr, T., Kreutzberg, G., Gemsa, D., Weber, E., & Moldofsky, H. (1995). A co-evolutionary theory of sleep. Med. Hypotheses. 45, 304–10. https://doi.org/10.1016/0306-9877(95)90122-1

Krueger, J., & Karnovsky, M. (1995). Sleep as a neuroimmune historical perspective phenomenon?: a brief historical perspective. Adv. Neuroimmunol. 5, 5–12. http://dx.doi.org/10.1016/0960-5428(94)00047-R

Krueger, J., & Majde, J. (1994). Microbial products and cytokines in sleep and fever regulation. Crit. Rev. Immunol. 14, 355–379. https://doi.org/10.1615/CritRevImmunol.v14.i3-4.70

Krueger, J., & Opp, M. (2016). Chapter Ten: Sleep and Microbes. International Review of Neurobiology. 131, 207–225. https://doi.org/10.1016/bs.irn.2016.07.003

Krueger, J., Obál, F., Fang, J., Kubota, T., & Taishi, P. (2001). The role of cytokines in physiological sleep regulation. Ann. N. Y. Acad. Sci. 933, 211–221. https://doi.org/10.1111/j.1749-6632.2001.tb05826.x

Krueger, J., Pappenheimer, J., & Karnovsky, M. (1982a). The composition of sleep-promoting factor isolated from human urine. J. Biol. Chem. 257, 1664–1669.

Krueger, J., Pappenheimer, J., & Karnovsky, M. (1982b). Sleep-promoting effects of muramyl peptides. Proc. Natl. Acad. Sci. U.S.A. 79, 6102–6106.

Krueger, J., Toth, L., Floyd, R., Fang, J., Kapás, L., Bredow, S., & Obál, F. (1994). Sleep, Microbes and Cytokines. Neuroimmunomodulation. 1, 100–109. https://doi.org/10.1159/000097142

Krueger, J., Walter, J., Dinarello, C., Wolff, S., & Chedid, L. (1984). Sleep-promoting effects of endogenous pyrogen (interleukin-1). Am. J. Physiol. 246, 994–999.

Krueger, J., Walter, J., & Levin, C. (1985). Factor S and related somnogens: an immune theory for slow-wave sleep. Brain. Mech. Sleep. 7, 253–276.

Kumar, A., Vashist, A., Kumar, P., Kalonia, H., & Mishra, J. (2012). Potential role of licofelone, minocycline and their combination against chronic fatigue stress induced behavioral, biochemical and mitochondrial alterations in mice. Pharmacol. Rep. 64, 1105–1115. https://doi.org/10.1016/S1734-1140(12)70907-6/S1734-1140(12)70907-6

Kushikata, T., Fang, J., & Krueger, J. (1999). Interleukin-10 inhibits spontaneous sleep in rabbits. J. Interferon. Cytokine. Res. 19, 1025–1030. https://doi.org/10.1089/ 107999099313244

Kushikata, T., Fang, J., Wang, Y., & Krueger, J. (1998). Interleukin-4 inhibits spontaneous sleep in rabbits. Am. J. Physiol. Regul. Integr. Comp. Physiol. 275, R1185–R1191.

Labus, J., Hollister, E., Jacobs, J., Kirbach, K., Oezguen, N., Gupta, A., Acosta, J., Luna, R., Aagaard, K., Versalovic, J., Savidge, T., Hsiao, E., Tillisch, K., & Mayer, E. (2017). Differences in gut microbial composition correlate with regional brain volumes in irritable bowel syndrome. Microbiome. 5, 49. https://doi.org/10.1186/s40168-017-0260-z

Lancel, M., Mathias, S., Faulhaber, J., & Schiffelholz, T. (1996). Effect of interleukin-1 beta on EEG power density during sleep depends on circadian phase. Am. J. Physiol. Regul. Integr. Comp. Physiol. 270, R830–R837.

Langille, M., Meehan, C., Koenig, J., Dhanani, A., Rose, R., Howlett, S., & Beiko, R. (2014). Microbial shifts in the aging mouse gut. Microbiome. 2, 50. https://doi.org/10.1186/s40168-014-0050-9

Leone, V., Gibbons, S., Martinez, K., Hutchison, A., Huang, E., Cham, C., Pierre, J., Heneghan, A., Nadimpalli, A., Hubert, N., Zale, E., Wang, Y., Huang, Y., Theriault, B., Dinner, A., Musch, M., Kudsk, K., Prendergast, B., Gilbert, J., & Chang, E. (2015). Effects of diurnal variation of gut microbes and high-fat feeding on host circadian clock function and metabolism. Cell. Host. Microbe. 17, 681–689. https://doi.org/10.1016/j.chom.2015.03.006

Louveau, A., Harris, T., & Kipnis, J. (2015). Revisiting the Mechanisms of CNS Immune Privilege. Trends. Immunol. 36, 569–577. https://doi.org/10.1016/j.it.2015.08.006

Masek, K., Kadlecová, O., & Petrovický, P. (1975). The effect of some bacterial products on temperature and sleep in rat. Z. Immunitatsforsch. Exp. Klin. Immuno. 149, 273–82.

Masek, K., Kadlecova, O., & Petrovicky, P. (1978). Pharmacological activity of bacterial peptidoglycan: the effect on temperature and sleep in the rat. Toxins. 1, 991– 1003. https://doi.org/10.1016/B978-0-08-022640-8.50094-90-08-022640-8.50094-9

Mašek, K., Kadlecová, O., & Rašková, H. (1973). Brain amines in fever and sleep cycle changes caused by streptococcal mucopeptide. Neuropharmacology. 12, 1039–1047. https://doi.org/10.10161016/0028-3908(73)90048-8/0028-3908(73)90048-8

Mazmanian, S., Cui, H., Tzianabos, A., & Kasper, D. (2005). An immunomodulatory molecule of symbiotic bacteria directs maturation of the host immune system. Cell. 122, 107–118. https://doi.org/10.1016/j.cell.2005.05.007.1016/j.cell.2005.05.007

McShane, B., Galante, R., & Biber, M. (2012). Assessing REM sleep in mice using video data. Sleep. 35, 433–442. https://doi.org/10.5665/sleep.1712

Meltzer, L., Serpa, K., & Moos, W. (1989). Evaluation in rats of the somnogenic, pyrogenic, and central nervous system depressant effects of muramyl dipeptide. Psychopharmacology. 99, 103–8. https://doi.org/10.1007/BF00634462

Morgun, A., Dzutsev, A., Dong, X., Greer, R., Sexton, D., Ravel, J., Schuster, M., Hsiao, W., Matzinger, P., & Shulzhenko, N. (2015). Uncovering effects of antibiotics on the host and microbiota using transkingdom gene networks. Gut. 64, 1732–1743. http://dx.doi.org/10.1136/gutjnl-2014-308820

Morrow, J., Vikraman, S., Imeri, L., & Opp, M. (2008). Effects of serotonergic activation by 5-hydroxytryptophan on sleep and body temperature of C57BL/6J and interleukin-6-deficient mice are dose and time related. Sleep. 31, 21–33. https://doi.org/10.1093/sleep/31.1.21

Mukherji, A., Kobiita, A., Ye, T., & Chambon, P. (2013). Homeostasis in intestinal epithelium is orchestrated by the circadian clock and microbiota cues transduced by TLRs. Cell. 153, 741–743. https://doi.org/10.1016/j.cell.2013.04.020

Nishino, R., Mikami, K., Takahashi, H., Tomonaga, S., Furuse, M., Hiramoto, T., Aiba, Y., Koga, Y., & Sudo, N. (2013). Commensal microbiota modulate murine behaviors in a strictly contamination-free environment confirmed by culture-based methods. Neurogastroenterol. Motil. 25, 521–528. https://doi.org/10.1111/nmo.12110

Nonaka, K., Nakazawa, Y., & Kotorii, T. (1983). Effects of antibiotics, minocycline and ampicillin, on human sleep. Brain. Res. 288, 253–259. https://doi.org/10.1016/0006-8993(83)90101-4

O’Mahony, S., Clarke, G., Borre, Y., Dinan, T., & Cryan, J. (2015). Serotonin, Tryptophan Metabolism and the Brain-Gut-Microbiome Axis. Behav. Brain. Res. 277, 32–48. https://doi.org/10.1016/j.bbr.2014.07.0272014.07.027

Opp, M. (2009). Sleeping to fuel the immune system: mammalian sleep and resistance to parasites. BMC Evolutionary Biology. 9, 8. https://doi.org/10.1186/1471-2148-9-8

Opp, M., & Imeri, L. (1999). Sleep as a behavioral model of neuro-immune interactions. Acta. Neurobiol. Exp. 59, 45–53.

Opp, M., & Krueger, J. (2017). Principles and practice of sleep medicine: chapter 19: sleep and host defense. 6, 193–201.e5. Philadelphia: Elsevier. https://doi.org/10.1016/B978-0-323-24288-2.00019-2

Oyama, N., Sudo, N., Sogawa, H., & Kubo, C. (2001). Antibiotic use during infancy promotes a shift in the TH1/TH2 balance toward TH2-dominant immunity in mice. J. Allergy. Clin. Immunol. 107, 153–159. https://doi.org/10.1067/mai.2001.111142

Pack, A., Galante, R., Maislin, G., Cater, J., Metaxas, D., Lu, S., Zhang, L., Von Smith, R., Kay, T., Lian, J., Svenson, K., & Peters, L. (2007). Novel method for high-throughput phenotyping of sleep in mice. Physiol. Genomics. 28, 232–238. https://doi.org/10.1152/physiolgenomics.00139.2006

Pappenheimer, J., Koski, G., Fencl, V., Karnovsky, M., & Krueger, J. (1975). Extraction of sleep-promoting factor S from cerebrospinal fluid and from brains of sleep-deprived animals. J. Neurophysiol. 38, 1299–1311.

Perlis, M., Kennedy, H., Salamone, C., Jungquist, G., Kochersberger, K., Plotkin, J., Allen, S., Karan, D., Ward, D., & Turnullo, S. (2006). Antibiotics may be insomnogenic. Poster presented at: Amer. Acad. Sleep. Med. 29, A259–A260.

Polanski, M., Johnson, T., & Karnovsky, M. (1992). Muramyl peptides as paradigms in neuroimmunomodulation. Ann. N. Y. Acad. Sci. 650, 218–220. https://doi.org/10.1111/j.1749-6632.1992.tb49125.x

Raper, D., Louveau, A., & Kipnis, J. (2016). How Do Meningeal Lymphatic Vessels Drain the CNS? Trends. Neurosci. 39, 581–586. https://doi.org/10.1016/j.tins.2016.07.001

Rea, K., Dinan, T., & Cryan, J. (2015). The microbiome: a key regulator of stress and neuroinflammation. Neurobiol. Stress. 4, 23–33. https://doi.org/10.1016/j.ynstr.2016.03.001

Reynolds, A., Broussard, J., Paterson, J., Wright, K., & Ferguson, S. (2017). Sleepy, circadian disrupted and sick: could intestinal microbiota play an important role in shift worker health? Mol. Metab. 6, 12–13. https://doi.org/10.1016/j.molmet.2016.11.004

Rhee, Y., & Kim, H. (1987). The correlation between sleeping-time and numerical change of intestinal normal flora in psychiatric insomnia patients. Bull. Nat. Sci. Chungbuk Natl. Univ. 1, 159–172.

Rogers, P., & Croft, M. (1999). Peptide dose, affinity, and time of differentiation can contribute to the Th1/Th2 cytokine balance. J Immunol. 163, 1205–1213.

Root-Bernstein, R., & Westall, F. (1990). Serotonin binding sites. II. Muramyl peptide binds to serotonin binding sites on myelin basic protein, LHRH, and MSH-ACTH 4-10. Brain. Res. Bull. 25, 827–841. https://doi.org/10.1016/0361-9230(90)90178-3

Ševcík, J., & Mašek, K. (1999). Brief communication the interaction of immunomodulator muramyl dipeptide with peripheral 5-HT receptors: overview of the current state. Int. J. Immunopharmacol. 21, 227–232. https://doi.org/10.1016/S0192-0561(98)00079-4

Shuman, M., Demler, T., Trigoboff, E., & Opler, L. (2012). Hematologic impact of antibiotic administration on patients taking clozapine. Innov. Clin. Neurosci. 9, 18–30.

Storch, C., Höhne, A., Holsboer, F., & Ohl, F. (2004). Activity patterns as a correlate for sleep–wake behaviour in mice. J. Neurosci. Methods. 133, 173–179. https://doi.org/10.1016/j.jneumeth.2003.10.008

Suda, K., Hicks, L., Roberts, R., Hunkler, R., & Taylor, T. (2014). Trends and seasonal variation in outpatient antibiotic prescription rates in the united states, 2006 to 2010. Antimicrob. Agents. Chemother. 58, 2763–2766. https://doi.org/AAC.02239-13

Sudo, N., Chida, Y., Aiba, Y., Sonoda, J., Oyama, N., Yu, X., Kubo, C., & Koga, Y. (2004). Postnatal microbial colonization programs the hypothalamic – pituitary – adrenal system for stress response in mice. J. Physiol. 558, 263–275. https://doi.org/10.1113/jphysiol.2004.063388

Thaiss, C., Levy, M., Korem, T., Dohnalov, L., Shapiro, H., Jaitin, D., David, E., Winter, D., Gury-BenAri, M., Tatirovsky, E., Tuganbaev, T., Federici, S., Zmora, N., Zeevi, D., Dori-Bachashi, M., Pevsner-Fischer, M., Kartvelishvily, E., Brandis, A., Harmelin, A., Shibolet, O., Halpern, Z., Honda, K., Amit, I., Segal, E., & Elinav, E. (2016). Microbiota diurnal rhythmicity programs host transcriptome oscillations. Cell. 167, 1495–1510. https://doi.org/10.1016/j.cell.2016.11.003

Thompson, R., Roller, R., Mika, A., Greenwood, B., Knight, R., Chichlowski, M., Berg, B., & Fleshner, M. (2017). Dietary Prebiotics and Bioactive Milk Fractions Improve NREM Sleep, Enhance REM Sleep Rebound and Attenuate the Stress-Induced Decrease in Diurnal Temperature and Gut Microbial Alpha Diversity. Front. Behav. Neurosci. 10, 240. https://doi.org/10.3389/fnbeh.2016.00240

Toth, L., Tolley, E., & Krueger, J. (1993). Sleep as a prognostic indicator during infectious disease in rabbits. Proc. Soc. Exp. Biol. Med. 203, 179–192. https://doi.org/10.3181/00379727-203-43590

Van Twyver, H., Webb, W., Dube, M., & Zackheim, M. (1973). Effects of environmental and strain differences on EEG and behavioral measurement of sleep. Behav. Biol. 9, 105–110. https://doi.org/10.1016/S0091-6773(73)80173-7

Vazquez-Palacios, G., Retana-Marquez, S., Bonilla-Jaime, H., & Velazquez-Moctezuma, J. (2001). Further definition of the effect of corticosterone on the sleep-wake pattern in the male rat. Pharmacol. Biochem. Behav. 70, 305–310. https://doi.org/10.1016/S0091-3057(01)00620-7

Vuong, H., Yano, J., Fung, T., & Hsiao, E. (2017). The microbiome and host behavior. Annu. Rev. Neurosci. 40, 21–49. https://doi.org/10.1146/annurev-neuro-072116-031347

Wimmer, M., Rising, J., Galante, R., Wyner, A., Pack, A., & Abel, T. (2013). Aging in mice reduces the ability to sustain sleep/wake states. PLoS. ONE. 8, e81880. https://doi.org/10.1371/journal.pone.0081880

Xie, L., Kang, H., Xu, Q., Chen, M. J., Liao, Y., Thiyagarajan, M., O’Donnell, K., Christensen, D., Nicholson, C., Iliff, J., Takano, T., Rashid, D., & Nedergaard, M. (2013). Sleep drives metabolite clearance from the adult brain. Science. 342, 373–377. https://doi.org/10.1126/science.1241224

Yaghouby, F., Donohue, K., O’Hara, B., & Sunderam, S. (2016). Noninvasive dissection of mouse sleep using a piezoelectric motion sensor. J. Neurosci. Methods. 259, 90–100. https://doi.org/10.1016/j.jneumeth.2015.11.004

Yano, J., Yu, K., Donaldson, G., Shastri, G., Ann, P., Ma, L., Nagler, C., Ismagilov, R., Mazmanian, S., & Hsiao, E. (2015). Indigenous bacteria from the gut microbiota regulate host serotonin biosynthesis. Cell. 161, 264–276. https://doi.org/10.1016/j.cell.2015.02.047

Zarrinpar, A., Chaix, A., Yooseph, S., & Panda, S. (2014). Diet and feeding pattern affect the diurnal dynamics of the gut microbiome. Cell Metab. 20, 1006–1017. https://doi.org/10.1016/j.cmet.2014.11.008

Zielinski, M., & Krueger, J. (2011). Sleep and innate immunity. Front. Biosci. 3, 632–642. http://dx.doi.org/10.2741/176

